# An agent-based model of molecular aggregation at the cell membrane

**DOI:** 10.1101/774505

**Authors:** Juliette Griffié, Ruby Peters, Dylan M. Owen

## Abstract

Molecular clustering at the plasma membrane has long been identified as a key process and is associated with regulating signalling pathways across cell types. Recent advances in microscopy, in particular the rise of super-resolution, have allowed the experimental observation of nanoscale molecular clusters in the plasma membrane. However, modelling approaches capable of recapitulating these observations are in their infancy, partly because of the extremely complex array of biophysical factors which influence molecular distributions and dynamics in the plasma membrane. We propose here a highly abstracted approach: an agent-based model dedicated to the study of molecular aggregation at the plasma membrane. We show that when molecules are modelled as though they can act (diffuse) in a manner which is influenced by their molecular neighbourhood, many of the distributions observed in cells can be recapitulated, even though such sensing and response is not possible for real membrane molecules. As such, agent-based offers a unique platform which may lead to a new understanding of how molecular clustering in extremely complex molecular environments can be abstracted, simulated and interpreted using simple rules.

**Author summary:** Molecular aggregation in cell membranes is a key component of cellular machinery, involved across cell types in inter-cellular communication and signalling pathway initiation. As such, understanding the underlying mechanisms and molecule cluster characteristics at a more theoretical level is a pre-requisite. Complete descriptive molecular models have proven impossible to realise due to the overall complexity of the processes involved, highlighting the need for novel approaches. While conceptual models have been shown to be powerful tools and are routinely used in other fields with high level of complexity such as social sciences or economics, they are overall lacking from the literature when it comes to cell studies. We suggest in this work that the same principle applies to cell biology and in particular, the study of molecular clustering. We propose here a general model, independent of cell types or signalling pathways: an agent-based model dedicated to molecular clustering in the plasma membrane. We show we are able to recapitulate molecular aggregation similar to observations in cells while new properties are highlighted by our model, for instance, clustering is a digitised process.

## Introduction

The plasma membrane (PM) has long been identified as a key component of cellular machinery. Its function is not limited to separating intra-cellular organelles from the extracellular environment; it is also recognized broadly as the principal instigator and coordinator of inter-cellular interactions[1,2]. The PM has been shown, for instance, to be involved in signalling pathway activation and inter-cell communication[3-5]. Processes such as neuronal transmission[6,7], the triggering of an immune response[8-10], Ras regulated pathways associated with mammalian cell survival and proliferation[11-13] and others, rely heavily on the molecular organisation at the PM.

In practice, molecular clustering has been highlighted as important in regulating signal transduction at the PM. Its study has benefited considerably from the rise of fluorescence microscopy and more recently super resolution methods[14-16], enabling, for the first time, the visualisation of molecular aggregation at the nanoscale. In fact, relatively small changes in the clustering configuration of signalling proteins, i.e. variations in their spatio-temporal organisation at the PM, are thought to be necessary and/or sufficient for the initiation of downstream signalling, a process commonly referred to as digitization[17-19].

Considering the current interest of the community in molecular aggregation at the PM, its understanding at a more theoretical level remains relatively understudied and simplified models are generally lacking or focusing on a specific activation pathway in a given cell type[20]. Agent-based models have been used across a wide variety of fields successfully to understanding collective behaviour, including in sociology[21], crowd management[22] and economics[23-25]. In this work, we develop an agent-based model dedicated to the study of molecular aggregation in the PM. The strength of such models with high levels of abstraction, such as the agent-based approach, is that they propose general rules otherwise inaccessible in complex systems. In our case, this translates to results being independent of cell type or signalling pathway. As such, the model presented here uniquely provides a platform in which one can study how to generate and modify molecular aggregation within a pre-set framework.

Our hypothesis is that all competing factors affecting a given molecular population distribution at the cellular membrane can be summarised as a single “desire for clustering” assigned to each agent (i.e. molecule). We demonstrate that this overall “desire for clustering” translated in an agent-based model, is sufficient to recapitulate clustering behaviours observed in cellular membranes. While in reality, molecules are not able to sense their neighbours at distances of tens or hundreds of nanometres, we show that they behave *as if* they can. We show that this apparent behaviour can be exploited to construct simplified models which recapitulate clustering profiles without the new to incorporate a full biophysical model. Interestingly, we also demonstrate that a modification of the environment, for instance a filamentous actin enriched environment, leads to distinct modifications of the clustering patterns as observed in cells.

## Model

### The theoretical framework

Any agent-based model relies on three levels of abstraction. It consists of an observer (or user) defining an environment (the PM in our case) in which agents (here, the diffusing molecules) evolve following pre-set rules. The model thus allows studying the fate of these agents over time (the third level of abstraction).

In practice, for simulations we consider 2000 agent-molecules diffusing freely (i.e. following Brownian motion) in a 2D 3 × 3 μm^2^ region of interest (ROI: the PM). Initially all proteins are randomly distributed forming a completely spatially random (CSR) distribution. The boundaries of the ROI are toroidally wrapped to ensure a constant number of molecules. The mobility characteristics of each molecule are updated every 10 ms. The simulations are run for 5 minutes (i.e. a total of 30,000 updated frames) as most cellular processes occur in this range of timescales. In all cases studied, n=30 simulations were generated.

Although a multitude of cellular processes have been shown to promote molecular clustering in cellular membranes, our initial hypothesis is that they can be summarised by a unique parameter: the “desire for clustering” given to each agent. This “desire” parameter results from a combination of competing cellular processes ranging from self-affinity, membrane lipid domains[26-32], the size exclusion model[33] and the picket-fence model[34] to name a few. It is this “desire for clustering” that constitutes the pre-set rules applied to each agent.

### Imposing pre-set rules on the agents: the “desire for clustering”

The method by which we generate a “desire for clustering” for each agent is by imposing a rate of Brownian motion, which is dependent on the immediate density of that agent’s local molecular environment. All agents possess a theoretical “target value” for clustering (defined as a certain number of neighbours within a 100 nm radius) and the closer its local density is to the targeted value, the slower the agent. Hence, whilst maintaining the assumption of molecules diffusing freely following Brownian motion in the PM, the diffusion coefficient (DC) of each molecule is fixed according to the local level of clustering.

For each frame and for each *i*^*th*^ localisation, the local density is estimated based on the “*L*_*R*_” value, i.e. the linearised localised Ripley’s K function:

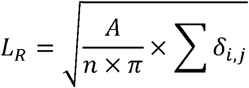

Where *A* is the area of the ROI, *n* is the total number of molecules in the ROI, for *i ≠ j, δ*_*i,j*_ = *1* if the molecule *j* is contained in a circle of radius *R* centred on molecule *i* and *δ*_*i,j*_ = *0* otherwise. Here, we define *R*=100 nm, as molecular clusters in cellular membranes have been shown to be typically below the diffraction limit of light (< 200nm)[12,35-37]. *L*_*R*_ is a mathematically well-defined quantity, considered as an accurate measure of local density and routinely used for spatial point pattern (SPP) cluster analysis [12,38-40]. For a CSR distribution, the average *L*_*R*_ value equals *R* (in our case *L*_*100*_ ∼ *100*, translating into 7 molecules encircled within 100 nm of each molecule). This is the starting point of the simulations **(Fig 1.a)**.

**Figure 1:**
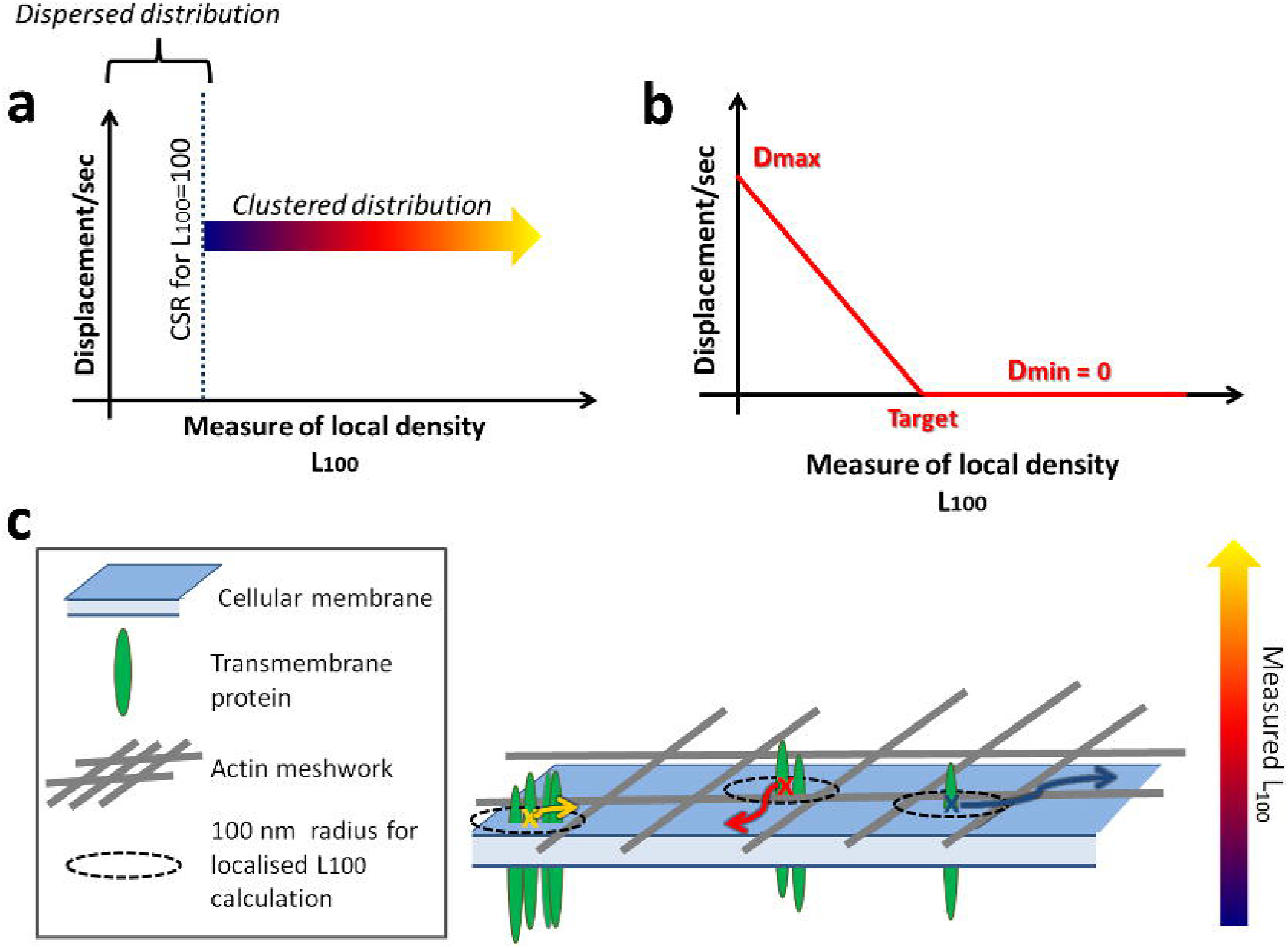
Defining the model. **a**. Defining the “desire for clustering” as a relation between the L100 value attributed to each molecule and its displacement per second. **b**. Displacement profile applied to each molecule in the Standard Condition. **c**. General model in which the displacement per second of each molecule (colour coded arrows) depends on the local level of clustering (i.e. the number of molecules encircled).

The target value for the clustering of all agents is simply defined as a specific value of *L*_*100*_. For each agent-molecule *i*, the measured value of *L*_*100,i*_ can therefore be compared to that target, from which the step displacement (*D*) per second is updated independently for each agent-molecule in every frame. Each molecule is thus attributed an independent D as a result of the level of clustering (*L*_*100*_), ranging from a pre-defined maximum (*D*_*max*_) to a minimum value when the agent reaches the targeted *L*_*100*_. Although the resulting diffusion is an approximation for Brownian motion, the difference in order of magnitude between step displacement (given the update/frame rate) and the length scale at which we study cluster formation, ensures this approximation is robust. Multiple functions can be chosen to map the dependence of *D* on *L*_*100*_. For the Standard Condition, we focus on a linearly converging profile in which a molecule *i* that has been attributed *L*_*100,i*_ >= *L*_*100,target*_ is set as fully immobile (i.e. D_min_ = 0) as illustrated in **Fig 1.b**. Our overall model is summarized in **Fig 1.c.**

After running the model, a number of outputs are generated. Principal among these are the average values of *L*_*100*_ per frame of the simulation, serving as a measure of the overall level of clustering within the ROI (referred to as 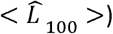). From the agent-molecule coordinates we can also extract more detailed clustering descriptors using a Bayesian based approach with topographic prominence thresholding[41-44]. These include the number of clusters, percentage of molecules in clusters, cluster radius and the number of molecules per cluster.

## Results

### The agent-based model leads to molecular aggregation

We first determined whether or not the “desire for clustering” is sufficient to induce measured clustering. For our Standard Condition displacement profile **(Fig 1.b)**, we fix Dmax at 3.5 nm.ms^−1^ which translates to a DC of ∼0.1 μm^2^ .s^−1^, i.e. within the range of expected values for transmembrane proteins (typically varying from 0.01 μm^2^.s^−1^ to 1 μm^2^.s^−1^)[45-47] and D_min_ = 0 (i.e. fully immobile). We varied the targeted *L*_*100*_ from 7 encircled molecules (*L*_*100*_ ∼ 100, the CSR case) up to 69 encircled molecules (*L*_*100*_ ∼ 317) with 2 molecule increments **(Fig 2.a)**. The range of targeted *L*_*100*_ values recapitulates clustering observed at the cellular membrane using single-molecule localisation, microscopy and quantitative cluster analysis techniques.

**Figure 2:**
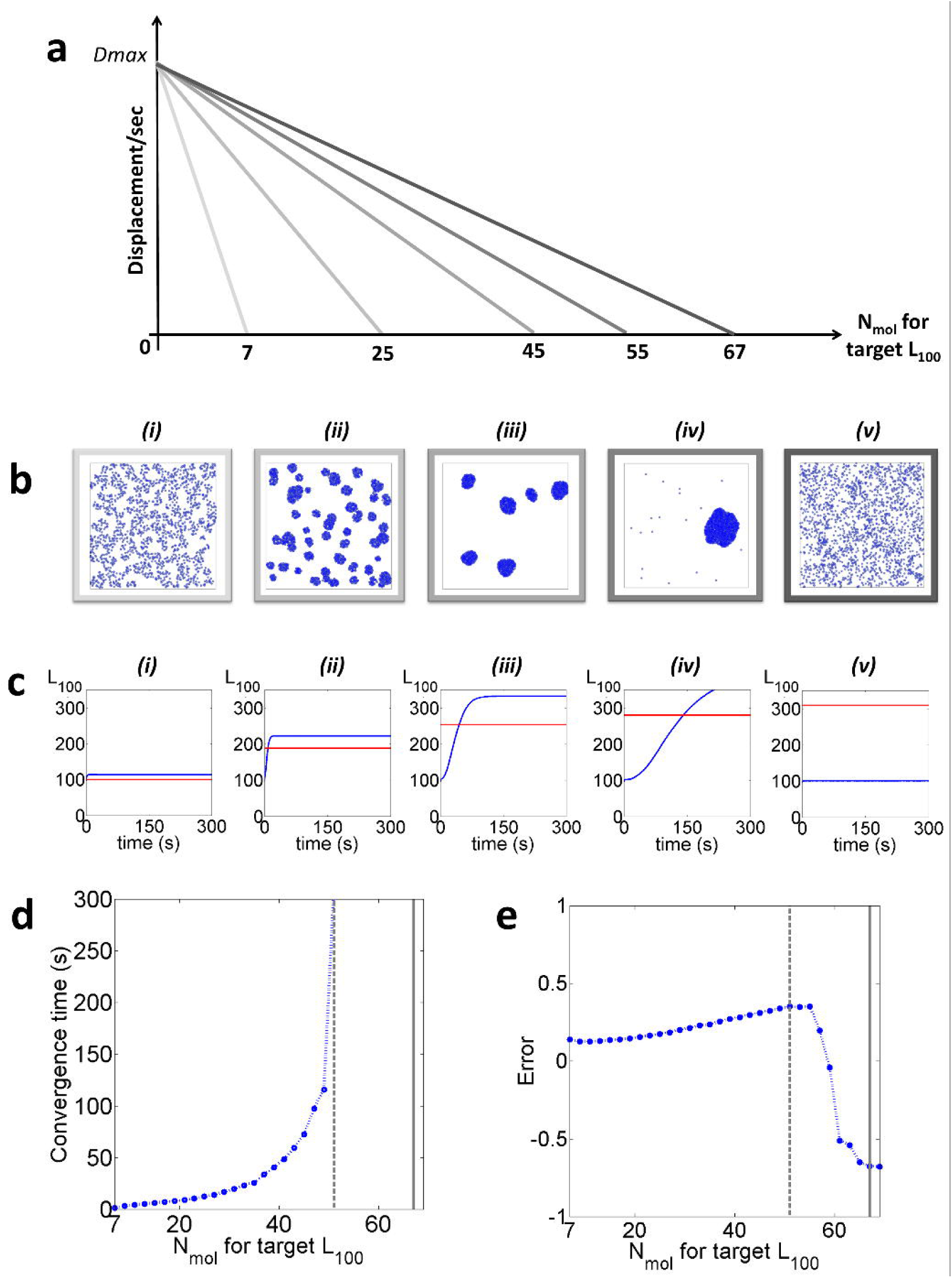
The Standard Condition (n=30 simulations per condition). **a**. Examples of displacement profiles applied to the agent-molecules. **b**. Examples of the molecular distributions obtained after 5 min simulations with the target values set at (i) 7, (ii) 25, (iii) 45, (iv) 55, (v) 67 encircled molecules. **c**. Corresponding 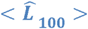 values extracted from all molecules over time (depicted blue) and their associated fixed targets (depicted red). d. Convergence time and e. Error as a function of the number of molecules for each *L*_*100*_ target (depicted in blue are the values obtained for each target, in dashed grey the point after which the system takes longer than 5 min to converge and in solid grey, the “cut-off” point).

**Fig 2.b** consists of representative examples of final frames (after 5 mins simulation) for 5 different target values (7, 25, 45, 55, 67 encircled molecules). They illustrate the agent-molecules reach a clustered distribution within 5 minutes simulations in most cases. These results demonstrate that the use of the “desire for clustering” as a summary of competing cellular processes is sufficient to induce molecular nanoscale aggregation, in turn supporting our initial hypothesis.

The 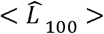 versus time plots, averaged over n=30 simulations per condition (**Fig 2.c**), confirm that different outcomes or “regimes” are indeed accessible, depending on the *L*_*100*_ target value applied. We can extract two instructive parameters from these curves: the convergence time (**Fig2.d**) and the associated error (**Fig 2.e**). The convergence time is defined as the amount of time it takes for 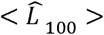 to reach a horizontal asymptote that is not equal to the initial CSR value. The error is calculated as: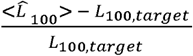. Both the convergence time and error highlight that 3 regimes are accessible within our timescale (**Fig 2.d-e**). Firstly, the case of full convergence: 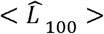 has reached a horizontal asymptote within the 5 minutes timescale (**Fig 2. b-c (i)-(iii)**). In this case the convergence time varies considerably from one target value to the next, increasing up to 49 encircled molecules, after which no asymptote is reached. The higher the target value is set, the longer the system takes to converge. In terms of error, this regime is associated with a systematic overshooting of the measured 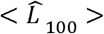 values compared to the target *L*_*100*_ value.

The second accessible regime (**Fig 2. b-c (iv)**) similarly consists of a converging system, but for which although displaying an increasing 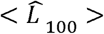 the horizontal asymptote has not been reached within the allocated 5 minutes timescale (between 51 and 65 encircled molecules).

The last accessible regime however differs drastically from previous cases. It consists of a non-converging system in which the agent-molecules continue to form a CSR distribution over 5 minutes whilst diffusing freely. Interestingly, the switch from converging regimes to CSR is a discontinuous digitised process with a cut-off point existing in the target *L*_*100*_value, after which the agent-molecule does not aggregate further (≥ 67 encircled molecules). We verified that this cut-off point was not in fact a slow convergence by repeating the simulations (target value = 69 encircled molecules) over a total 30 minutes, which demonstrates no sign of convergence (**Fig S1**).

**Fig 3** summarizes the associated cluster descriptors extracted from the last simulated frame, averaged over the n=30 simulations for every target *L*_*100*_ value condition. These results confirm the existence of 3 possible molecular distribution outcomes, which, in this case translates to: a fully clustered distribution, a clustered distribution with a small ratio of remaining unclustered molecules and finally, a CSR distribution after 5 minutes (**Fig 3.a**). Overall, these results confirm that in most cases the “desire for clustering” is a sufficient rule to induce agent-molecule aggregation. In practice, for the clustered cases the final distribution arises from the appearance of molecular “nucleation sites” in a matter of seconds (**Movie S1**). As in cells, an existing cluster appears to facilitate the aggregation of close by molecules[33,48].

**Figure 3:**
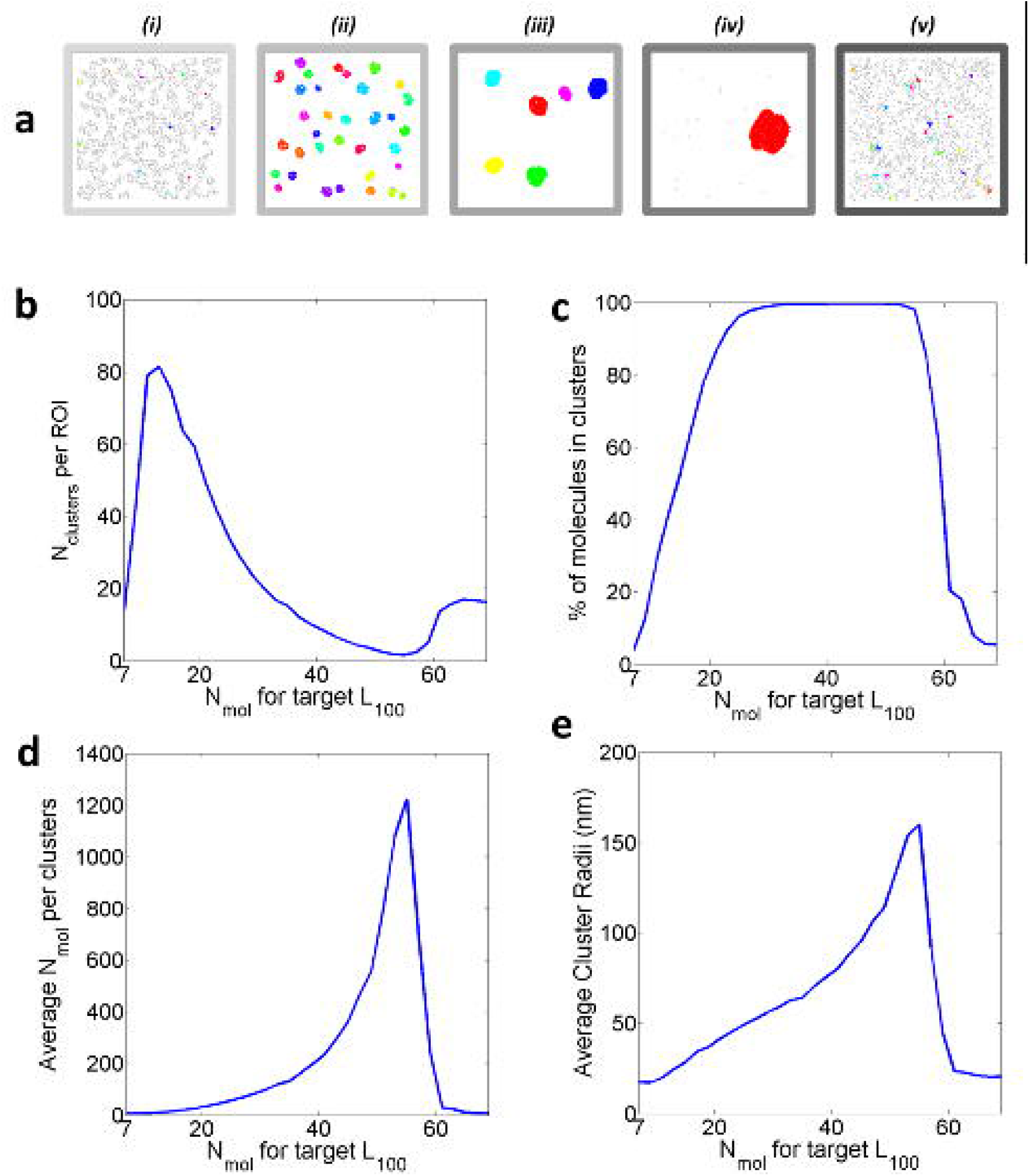
Standard Condition clustering characteristics. **a**. Example of analysed cluster maps pseudo coloured by cluster identification numbers generated from the final simulated frames with the target set to (i) 7, (ii) 25, (iii) 45, (iv) 55, (v) 67 encircled molecules. For the n=30 simulations: **b**. Average number of clusters per ROI, **c**. % of clustered molecules, **d**. Average number of molecules per clusters, **e**. Average cluster radius (nm).

Importantly, our results highlight that for subtle variations of the target value, the resulting molecular distribution present diverse clusters characteristics (**Fig 3.b-e**). We can generate, for instance, simulations with an average of 5.5 clusters of 95 nm radius with a target set at 45 encircled molecules, whilst a target set at 49 leads to only 3.6 clusters of 114 nm radius. As such, our agent-based model compliments phenomena observed in cells, for which very small changes, for example the engagement of a receptor protein on the PM or the modification of membrane tension are associated with rapid and dramatic changes in molecular clustering. Overall, the clustered distributions obtained via simulation resemble clusters found in cells (their radii varying from 24.5 to 160.2 nm, for instance). These results again verify our initial hypothesis, and more broadly, the validity of the agent-based model presented here for the study molecular aggregation in the PM. As it was the case for the study of the 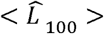 time-series curves (**Fig 2c**), detailed cluster descriptors also display a digitized profile with a cut-off point after which the molecules remain in a CSR distribution. This data therefore suggests that clustering itself could be a digitized process with as precise band pass. We further investigated the role of R on the outcome, studying the impact of smaller radius (R=50nm in **Fig S2**) and radius above the diffraction limit (R=200nm in **Fig S3**). In all cases, the three available regimes previously observed (converged, converging, unconverging) are robust to variation in R. Interestingly, the cut-off point appears to be function of R.

### Modifying the pre-set rules applied to the agent-molecules

We further study the impact of the pre-set rules applied to the agent-molecules on the measured clustering outcomes. For that purpose, we first varied the parameter *D*_*max*_ between a fast and low diffusion (respectively corresponding to DC∼1 μm^2^.s^−1^ and 0.01 μm^2^.s^−1^) (**Fig 4.a**). Interestingly, the digitised behaviour, as well as the overall trends of both the 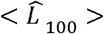 curves and the cluster descriptors initially observed in the Standard Condition, are maintained for both the fast and slow condition. The cut-off value for convergence however appears to be highly dependent on the *D*_*max*_ value used for the displacement curves (target value = 59 and 69 encircled molecules for the fast and slow case respectively) (**Fig 4.b-c**), with slower diffusion leading to a thinner pass band for which the system converges to a clustered distribution.

**Figure 4:**
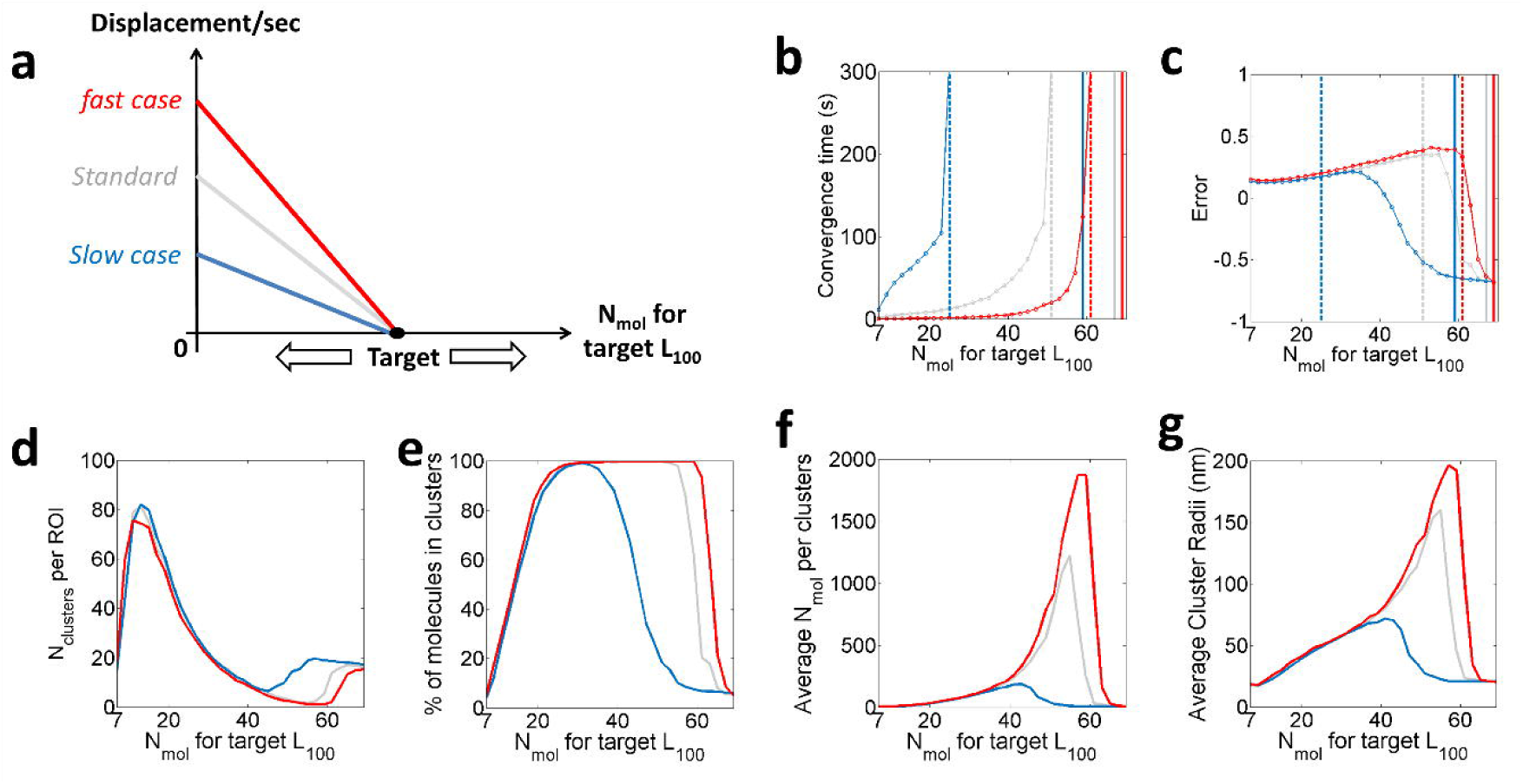
Varying Dmax for n=30 simulations per condition. In red the fast case (∼ 1 μm.s^−1^), in grey the Standard Condition (∼ 0.1 μm.s^−1^) and in dark blue the slow case (∼ 0.01 μm.s^−1^). **a**. Illustration of the speed profiles applied to the agent-molecules. **b**. Convergence time for the 3 cases, the target after which the system takes longer than 5 minutes to converge (dashed vertical line), and the “cut-off” point (solid vertical line). **c**. Error for each of the 3 D_max_ conditions. **d**. The average number of clusters per ROI. **e**. The % of clustered molecules. **f**. The average number of molecules per clusters. **g**. The average cluster radius (nm), extracted from the final molecular distribution.

In particular, these results suggest that faster diffusing molecules tend to aggregate more quickly (i.e. lower convergence time) and over a larger range of target values. As they travel larger distances, fast moving molecules are more likely to be caught within the range of influence of a spontaneous nucleation site (**Movie S2 and S3**). For fast diffusing molecules, the associated simulations converge to clustered distributions even for *L*_*100*_ targets which do not converge for slower diffusing molecules, whilst forming larger aggregates (**Fig 4.d-g**). For instance, for a target set at 63 encircled molecules, the fast diffusing model achieves an average of 7.6 clusters containing an average of 157 molecules while this target remains unclustered for the slow case. Again, our model recapitulates phenomena observed in cells for which faster diffusing molecules result in clustered distribution differing from slowly diffusing molecules. Decreased membrane viscosity, associated with faster diffusing transmembrane proteins, for instance, has been shown to result in more clustered distribution[49].

So far, we have chosen to vary the displacement (or per frame step size) for each molecule over time, which is a robust approximation of Brownian motion at the length scale at which we are studying cluster emergence versus frame update. However, a more general approximation consists in a random walk in which the per-frame step sizes are drawn randomly (and independently) from a normal distribution centred on the targeted diffusion coefficient. Using this approach, we were able to show that not only molecular aggregation can be achieved but also that similar mode (converged, converging, unconverging) are observable (Fig S4).

We also studied the impact of the shape of displacement curves themselves on the output structures (**Fig S5**). Surprisingly, in both non converging-cases, the three possible outcome regimes (converged, converging, unconverging) remain, with similar values obtained for the cut-off points. This suggests that the digitisation of molecular aggregation is a phenomenon independent of the displacement-clustering relation rule imposed on the agent-molecules. However, the nanoscale structures obtained in both cases differ drastically from the Standard Condition. We further demonstrate that both pores and mobile clusters are achievable within our agent-based model (**Movie S4 and S5**). Importantly, both molecular pores[50] and dynamic clusters[51,52] exist in cellular membranes, highlighting the applicability of our approach to a vast range of molecular nanoscale structures and processes.

We further investigated the impact of D_min_ for non-converging cases. We set D_min_ as the expected per frame displacement of a cluster containing all molecules in the ROI (**Fig S6**), 20% of the molecules (**Fig S7**) and 10% of the molecules (**Fig S8**). In all cases, although, we observe a conservation of the three possible outcomes, the cut-off point itself appears to be highly dependent on D_min_.

Finally, whilst the use of a linear decay (whether converging or not), is necessary to understand better the mechanism behind the emergence of molecular aggregation, it is unlikely to recapitulate all the variations existing at the cell membrane. We run additional simulations with a desire for clustering following a non-linear decay (quadratic) to verify the robustness of clusters emergence. As in the linear case, we observe molecular aggregation (**Fig S9**).

### The impact of the actin mesh on molecular aggregation

One of the advantages of performing simulations such as the agent-based model presented here, is that it enables studying the impact of any modifications in the environment architecture on the agents: e.g. would agent behaviour be different in the presence of obstacles or boundaries within the ROI. This approach is particularly interesting in the case of the cellular membrane whose nanoscale architecture is well characterised. Our framework can therefore be used to understand the impact of known cellular structures on the basal level molecular aggregation determined only by the desire for clustering. Here, we focus on the role played by the actin meshwork situated directly beneath the PM. In particular, we compare our Standard Condition with an actin enriched environment. Certainly, as previously stated, the desire for clustering encompasses complex, intertwined and diverse cellular processes, including the potential role of the actin mesh. However, here we are able to study the effect of an increase in filamentous actin added to the desire for clustering as defined in our Standard Condition or basal level of clustering.

Actin is a fibrous polymer protein which forms a key component of the cell skeleton. It allows the cell to generate forces through actin polymerisation which serves as a highway for molecular trafficking by motor proteins and enhances the cells’ structural stability[53-56]. This final function is achieved by a dense meshwork of actin placed immediately below the PM (the actin cortex). Recently, research has focussed on how this cortical actin affects the diffusion and clustering of membrane proteins by essentially forming semi-permeable barriers proximal to the bilayer, transiently confining membrane proteins[57-59]. By modifying the density of this filamentous actin mesh, the enclosed areas of the PM in which molecules can diffuse freely varies. The fact that key signalling pathways from or to the PM are associated with substantial remodelling of the actin meshwork[60-62] tend to sustain these theories of the importance of the actin mesh in regulating molecular aggregation at the PM across multifarious cell types.

In practice, actin fibres are translated into our agent-based model as reflective boundaries arranged in a regular grid, within the ROI. We vary the meshwork density from relatively sparse (1000 nm spacing) to more dense cases (500 nm and 250 nm spacing). 7 conditions were simulated with: 7, 15, 25, 35, 45, 55 and 65 encircled molecules targets. Representative examples of the molecular distributions obtained after 5 mins simulations are illustrated in **Fig 5.a-c.** These results suggest that the system can still reach a clustered distribution despite the presence of the additional imposed actin meshwork. These results further highlight that the presence of added actin meshwork and more specifically its density, can modify both the clustering characteristics of the agents and the cut-off point.

**Figure 5:**
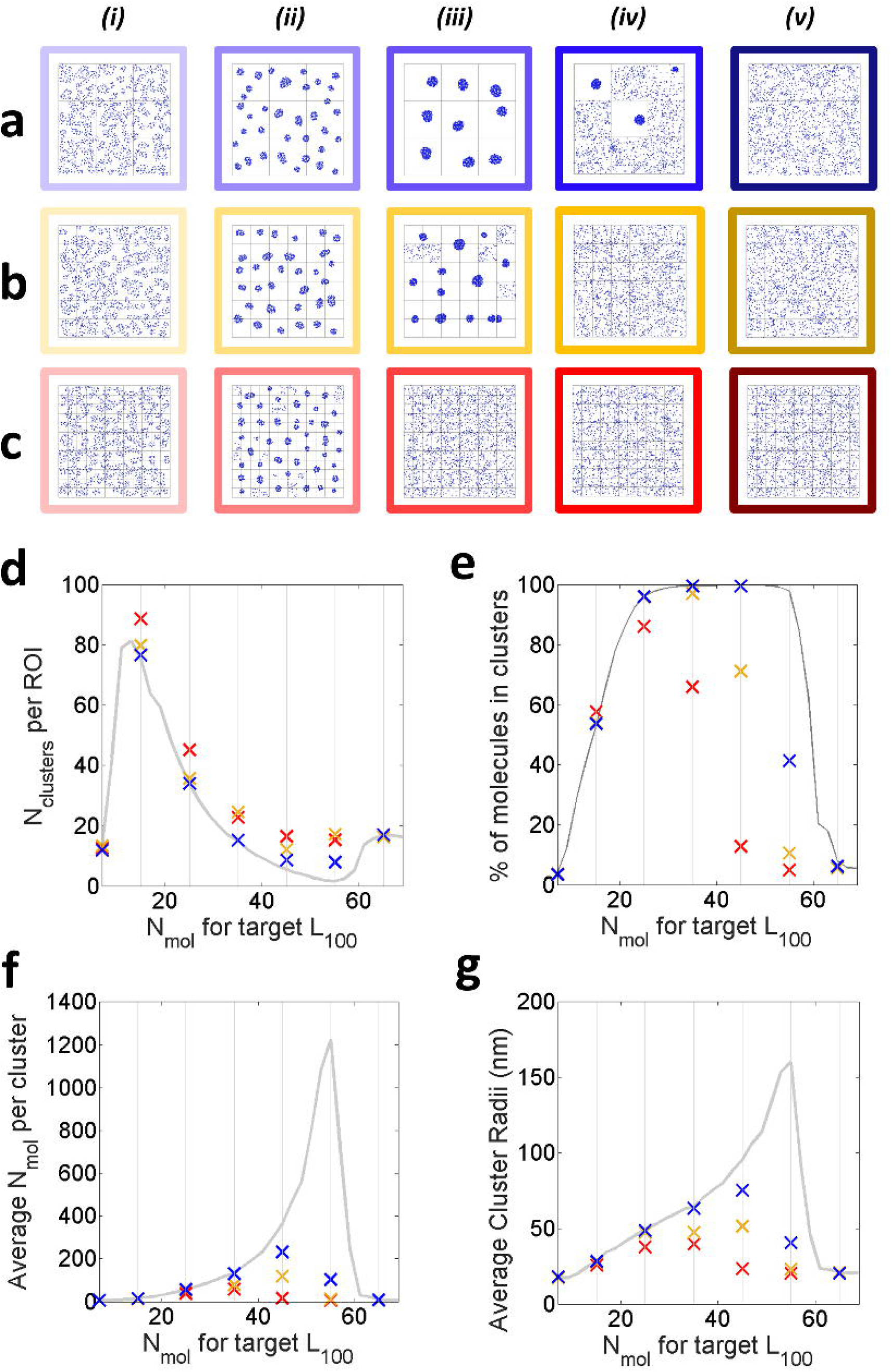
An actin enriched environment. **a**,**b**,**c**. Examples of the molecular distributions obtained after 5 mins simulations with the target encircled molecules set to (i) 7, (ii) 25, (iii) 45, (iv) 55, (v) 65 with 1000 nm, 500 nm and 250 nm spaced actin boundaries respectively. **d**. Average number of clusters per ROI, **e**. % of clustered molecules, **f**. Average number of molecules per clusters, **g**. Average cluster radius (nm). For the n=30 simulations, the 1000 nm, the 500 nm, 250nm cases are depicted blue, orange and red respectively (the grey curves represent the Standard Condition).

We studied the impact of filamentous actin on the overall 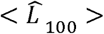 trends, similarly to observations made when varying the displacement profile starting point D_max_. Our simulations reach the cut-off point for a target value smaller than 55, 65 and 65 encircled molecules for 250 nm, 500 nm and 1000 nm mesh spacing respectively. In all cases, the cut-off values consist of smaller values than in the absence of a mesh (i.e. in the Standard Condition). However, although the digitised behaviour is preserved, with varying pass band width depending on the mesh density, our results also suggest that the actin mesh alone, in particular its density, impacts on the characteristics of the molecular aggregates for the same fixed target value. The presence of a dense actin mesh results in agent-molecules reaching a clustered distribution over a very limited range target values, whilst systematically forming more numerous smaller clusters when converging (**Fig 5. e-g**). For instance, the target value set at 25 encircled molecules in the Standard Condition results in an average of 34 clusters of 57 molecules, whereas in the case of an actin fibres enriched environment, we find an increase in the number of clusters (45.2, 35.7, 34.0 for 250, 500 and 1000nm spacing respectively) containing less molecules (38.1, 53.7, 56.5 for 250, 500 and 1000nm spacing respectively). Again, our results compliment phenomena expected with the picket fence model[34] in cells in which actin polymerisation and organisation in regions of dense meshwork is associated with more numerous but importantly, less dense clusters.

## Discussion

The cellular membrane has been shown to be a system with a high level of complexity: various processes are known to be involved in the regulation of molecular distributions at the PM. For the most part, these processes (e.g. the picket fence theory, protein affinity, the size exclusion model, the lipid raft hypothesis) are thought to act locally with often intertwined and competing effects on molecular aggregation. Their complex relationships remain to be fully characterised and more processes are likely to be identified with technological advancement in microscopy. In fact, there is no clear consensus in the field on the role of each in regards of the other[63], as well as strong controversies on the existence of some of these main players (i.e. lipid rafts)[47,64].

Because of these complex factors, complete and general molecular models, for example atomistic MD simulations of the whole plasma membrane-cortex system, have proven impossible to realise. We suggest in this work that cellular biology will benefit vastly from conceptual models such as agent-based approach as a way to improve our understanding of key cellular processes (e.g. protein clustering at the PM) similarly to other fields with high level of complexity like economics[25]. The agent-based model presented here may lead to simpler ways to study, simulate and interpret molecular clustering in the membrane without using complex biophysical models. We demonstrate that simple local rules obeyed by the agents, can lead to the emergence of aggregation at macroscale. More precisely, we show that the processes acting on molecules in cellular membranes can be summarized in a unique parameter: the “desire for clustering” and that through this framework, the molecular clustering observed in cells can be recapitulated. Our results also suggest that clustering itself is a digitized process in addition to clusters ability to digitise the flow of information along signalling pathways

Interestingly, our simulations highlight that for small variations of the “desire for clustering” applied to molecules, drastically different clustering outcomes are obtained. This mimics observations made in cells where small changes at the PM are associated with significant modifications of the molecular distributions, possibly facilitating signalling or effector functions[65]. Our results also confirm that other known impact of cellular processes on clustering can be recapitulated by the agent-based model presented here. For instance, we show that faster diffusing molecules tend to form more clustered distributions, while enrichment in filamentous actin compared to the basal level, impacts directly on the clusters descriptors with smaller yet more numerous clusters. Both phenomena have been previously observed in cells, demonstrating that the model can generate distributions consistent with those previously observed experimentally. Adding more complex environments, such as non-fully reflective mesh, in which the distribution of agents can evolve, could help further characterise the dependency of cluster emergence to these precise architectures.

Agent based models have particular advantages over other methods e.g. MD simulations, which make them particularly useful for applications studying membrane protein distributions and dynamics. Firstly, they do not require complete knowledge of the structure of the system under investigation. They are well suited to the study of emergent phenomena – where macroscale architecture is generated from underlying local forces and they provide a simple and intuitive conceptual framework for understanding how biophysical phenomena (e.g. actin) impact molecular behaviour. In that context, agent based models might be useful for the design of pharmacological interventions designed to alter molecular nanoscale organisation due to their ability to predict the macroscale outcome of a local perturbation. For example, new therapeutic concepts involve the modification of plasma membrane clustering in the context of disease[66].

## Acknowledgements

This work was supported by ERC Starter Grant #337187 to DMO.

## Author contributions

JG, DMO designed the model, JG implemented the model, JG developed and ran the simulations, JG, RP ran the analysis, JG, DMO wrote the manuscript.

