## Supplementary Information for "An agent-based model of molecular aggregation at the cell membrane"

### Supplementary figures

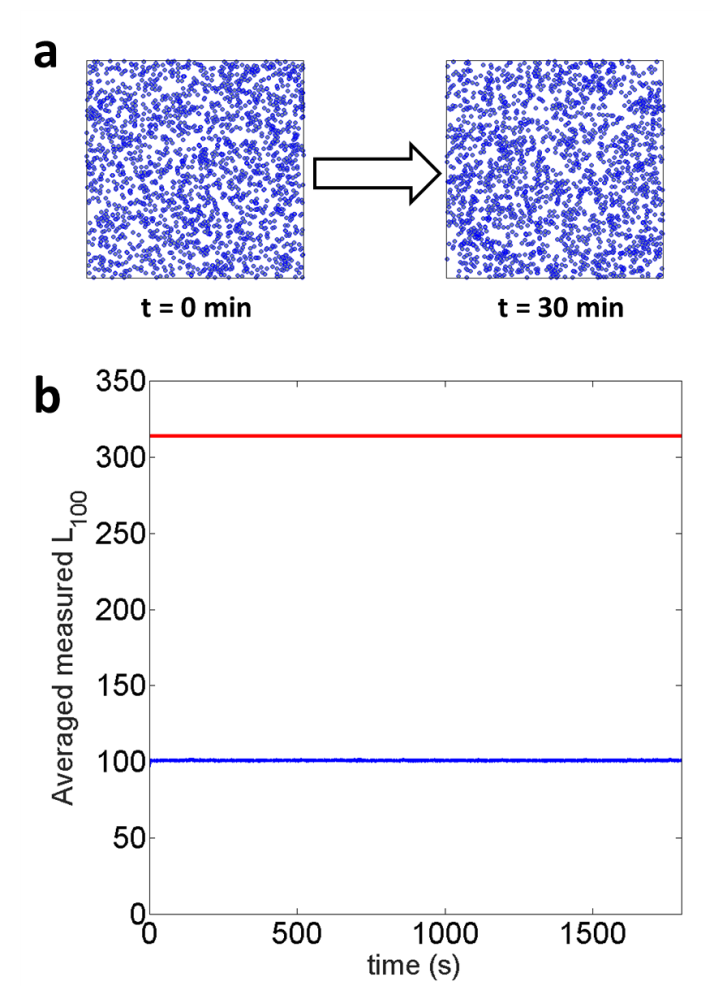

Figure Supplementary 1: 30 min simulation with the target value set at 69 encircled molecules in the Standard Condition, n=30. a Molecular distribution from t=0 min to t=30 min. b. Resulting $\boldsymbol{<}{\hat{\boldsymbol{L}}}_{\boldsymbol{100}}\boldsymbol{>}$ curve.

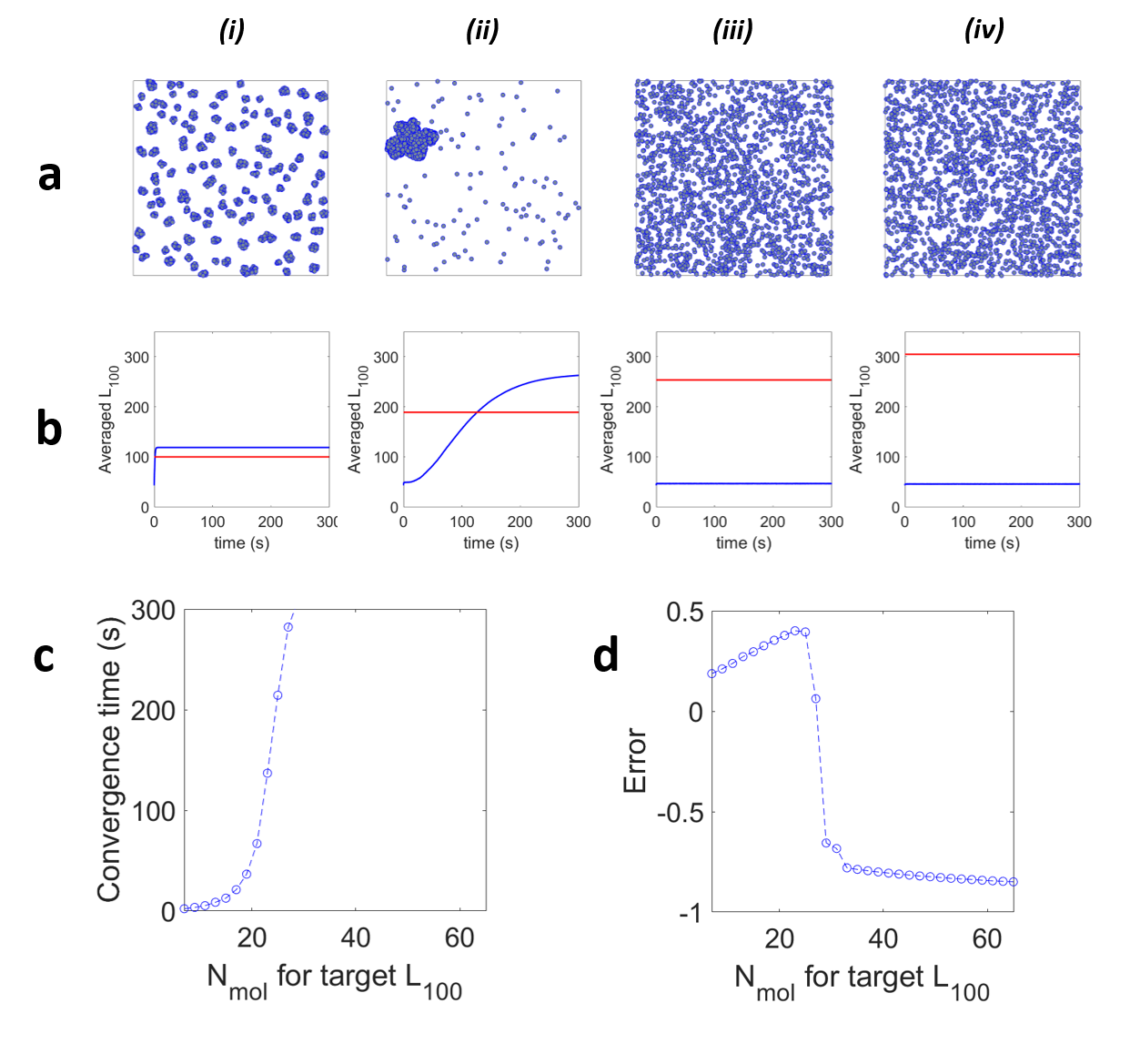

Figure Supplementary 2: Converging scenario with R=50nm, (n=30 simulations per condition). a. Examples of the molecular distributions obtained after 5 min simulations with the target values set at (i) 7, (ii) 25, (iii) 45, (iv) 55, (v) 65 encircled molecules. b. Corresponding $\boldsymbol{<}{\hat{\boldsymbol{L}}}_{\boldsymbol{100}}\boldsymbol{>}$ values extracted from all molecules over time (depicted blue) and their associated fixed targets (depicted red). c. Convergence time and d. Error as a function of the number of molecules for each *L_100_* target (depicted in blue are the values obtained for each target).

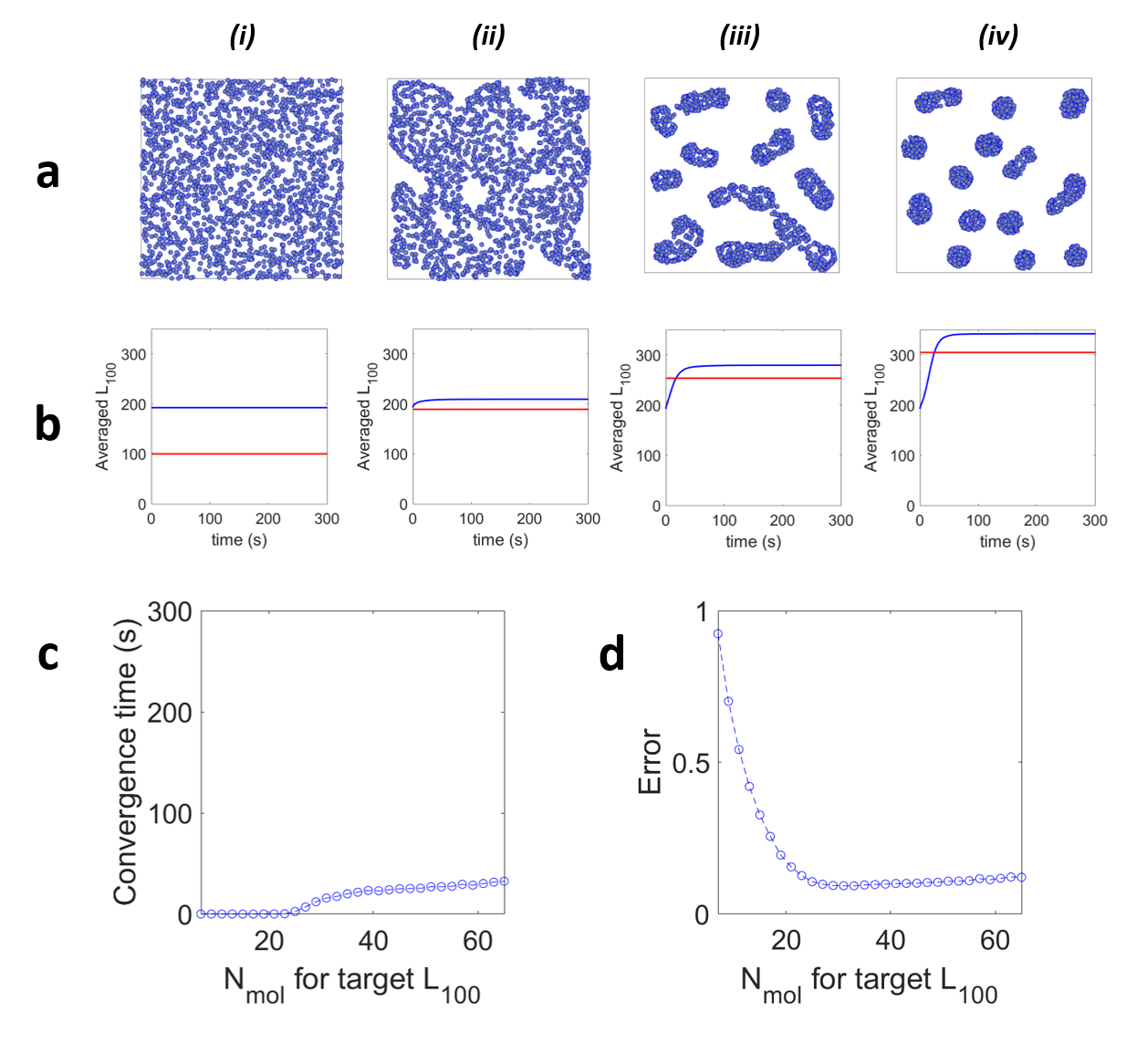

Figure Supplementary 3: Converging scenario with R=200nm, (n=30 simulations per condition). a. Examples of the molecular distributions obtained after 5 min simulations with the target values set at (i) 7, (ii) 25, (iii) 45, (iv) 55, (v) 65, (vi) 80, (vii) 90 encircled molecules. b. Corresponding $\boldsymbol{<}{\hat{\boldsymbol{L}}}_{\boldsymbol{100}}\boldsymbol{>}$ values extracted from all molecules over time (depicted blue) and their associated fixed targets (depicted red). c. Convergence time and d. Error as a function of the number of molecules for each *L_100_* target (depicted in blue are the values obtained for each target).

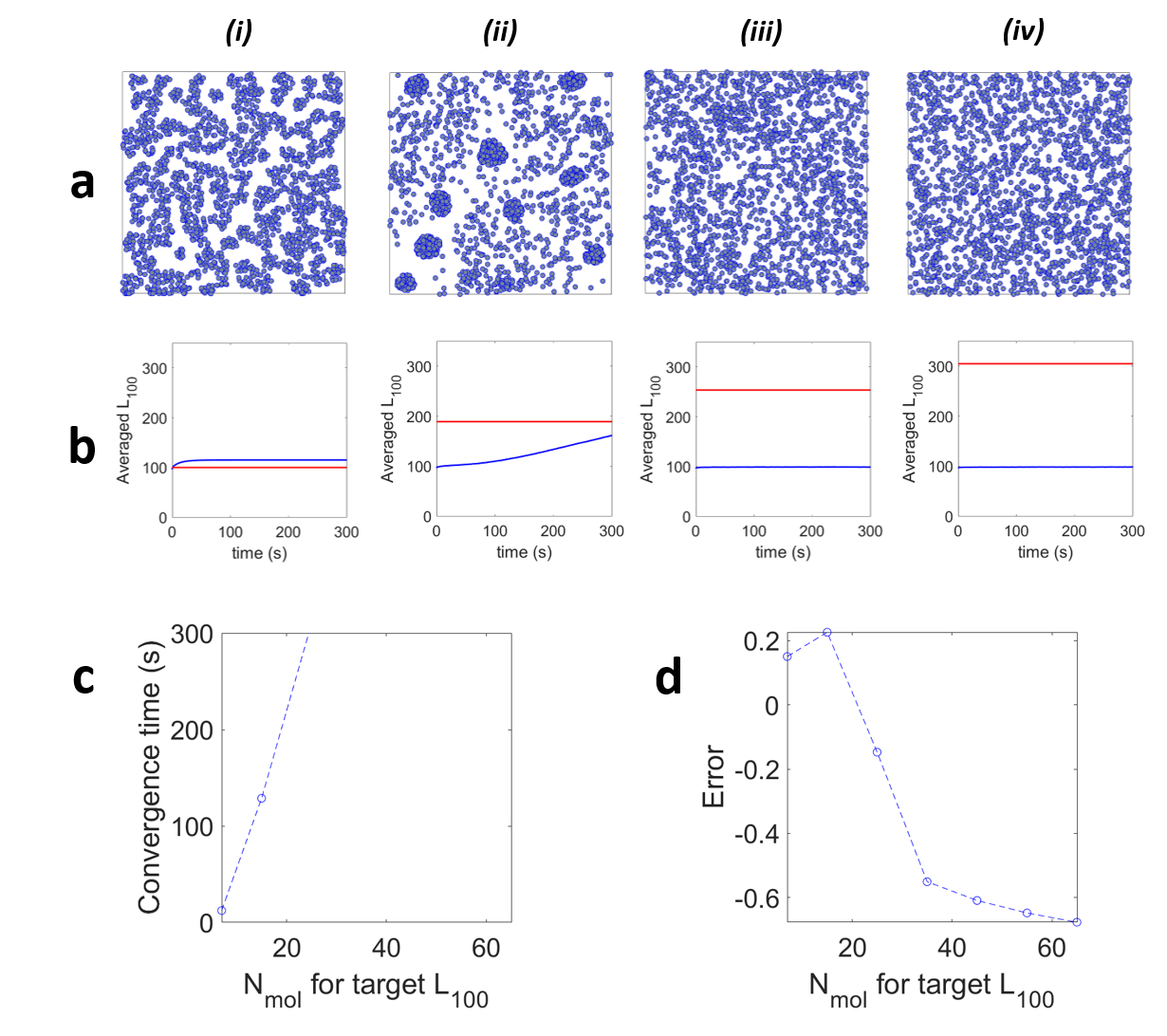

Figure Supplementary 4: Diffusion coefficient extracted from normal distributions, (n=30 simulations per condition). a. Examples of the molecular distributions obtained after 5 min simulations with the target values set at (i) 7, (ii) 25, (iii) 45, (iv) 55, (v) 65 encircled molecules. b. Corresponding $\boldsymbol{<}{\hat{\boldsymbol{L}}}_{\boldsymbol{100}}\boldsymbol{>}$ values extracted from all molecules over time (depicted blue) and their associated fixed targets (depicted red). c. Convergence time and d. Error as a function of the number of molecules for each *L_100_* target (depicted in blue are the values obtained for each target).

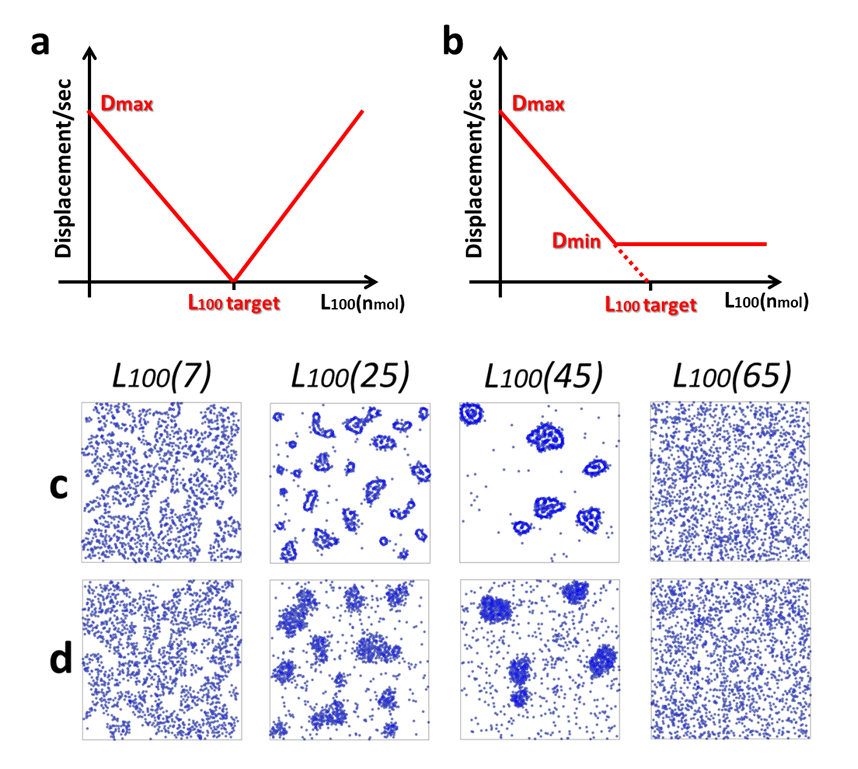

Figure Supplementary 5: Varying the displacement profile. a. Non-converging scenario where a molecule i is immobile for L100,i = L100,target only. b. Converging scenario with Dmin>0. c. Example of final updates frame for target set at 7, 25, 45, 65 encircled molecules, in the non-converging scenario. d. Example of final updates frame for target set at 7, 25, 45, 65 encircled molecules in the converging scenario with Dmin>0.

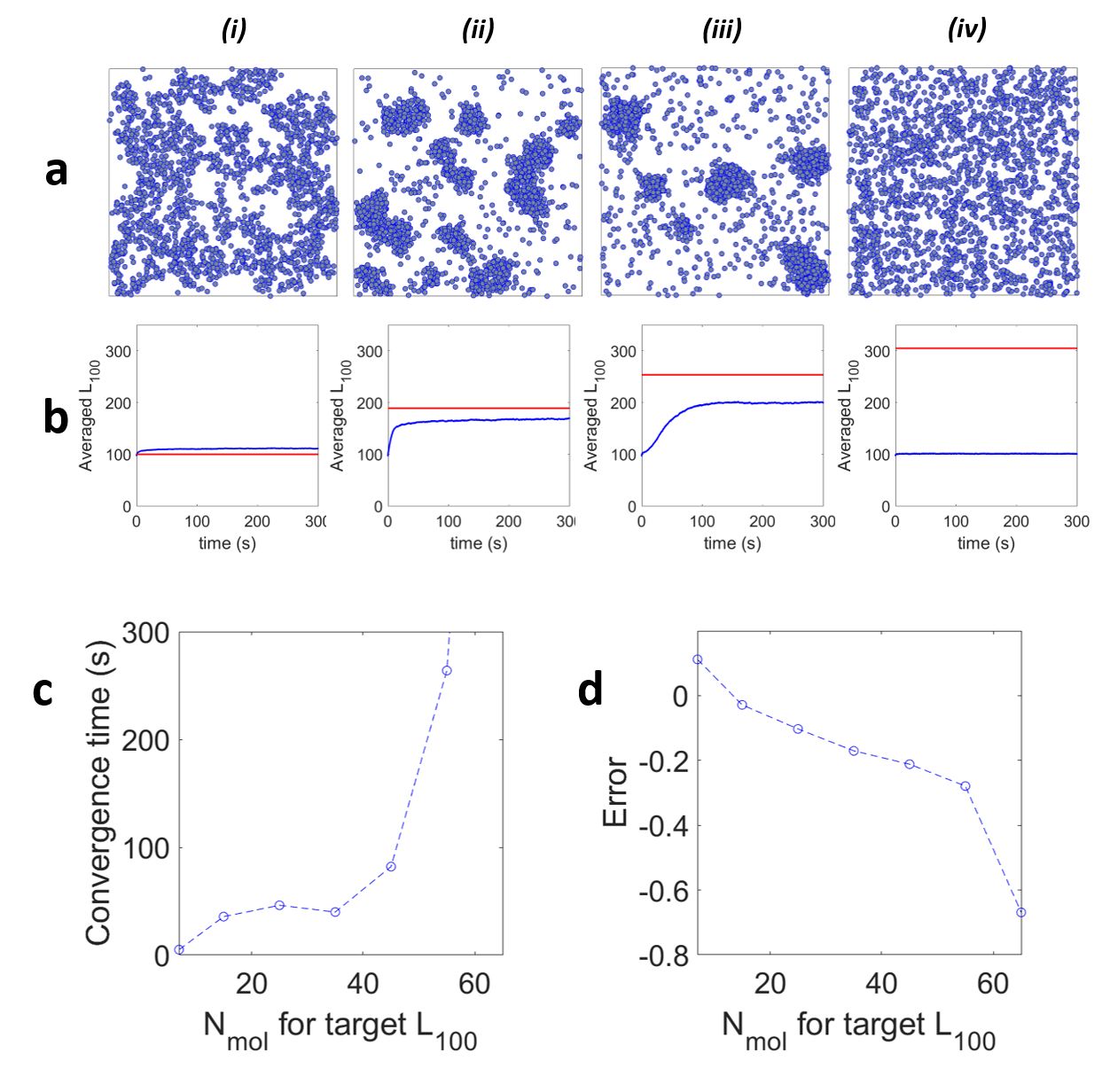

Figure Supplementary 6: Converging scenario with Dmin ~ 0.52 nm.ms^-1^, (n=10 simulations per condition). a. Examples of the molecular distributions obtained after 5 min simulations with the target values set at (i) 7, (ii) 25, (iii) 45, (iv) 55, (v) 65, encircled molecules. b. Corresponding $\boldsymbol{<}{\hat{\boldsymbol{L}}}_{\boldsymbol{100}}\boldsymbol{>}$ values extracted from all molecules over time (depicted blue) and their associated fixed targets (depicted red). c. Convergence time and d. Error as a function of the number of molecules for each *L_100_* target (depicted in blue are the values obtained for each target).

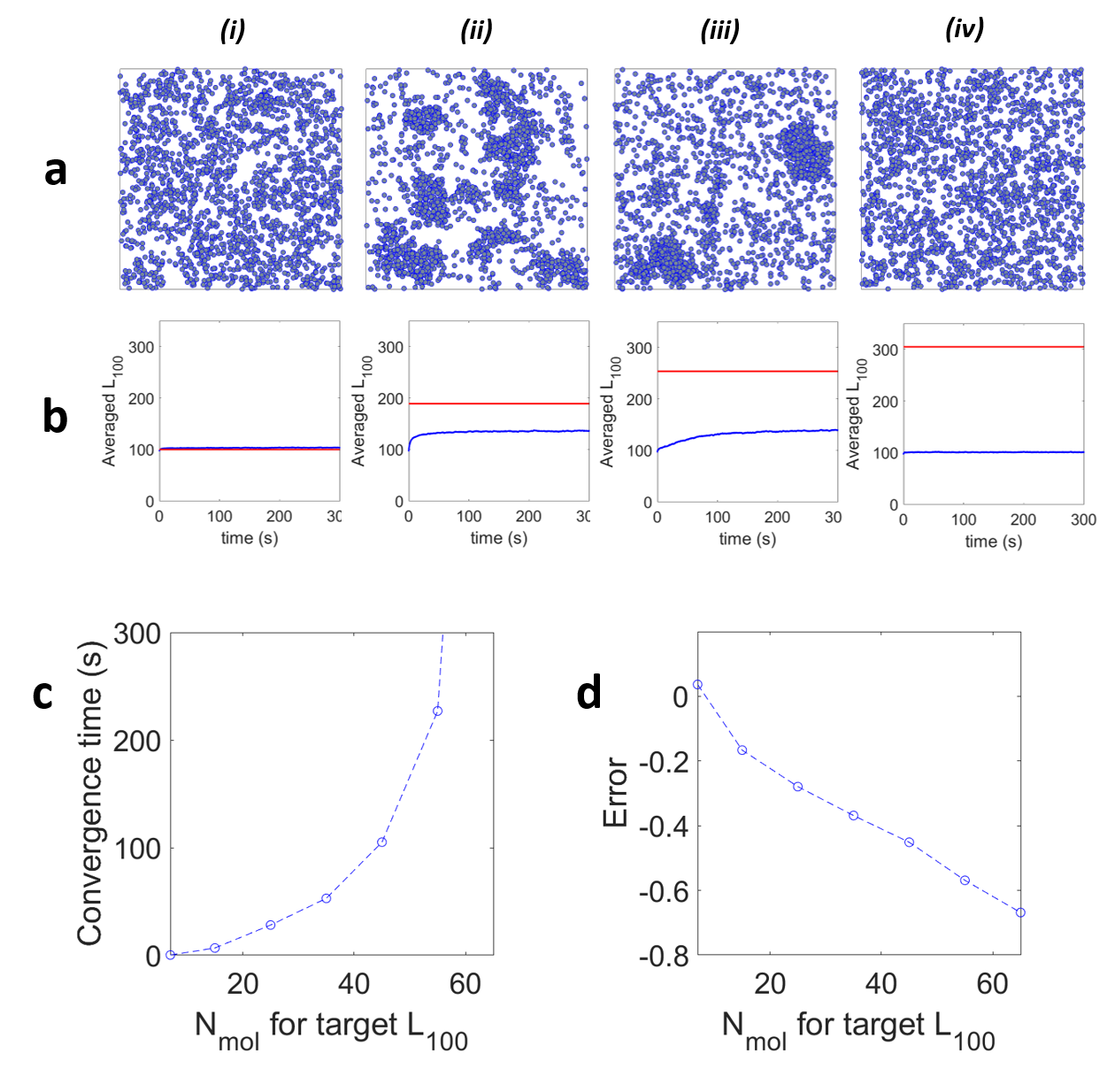

Figure Supplementary 7: Converging scenario with Dmin ~ 0.78 nm.ms^-1^, (n=10 simulations per condition). a. Examples of the molecular distributions obtained after 5 min simulations with the target values set at (i) 7, (ii) 25, (iii) 45, (iv) 55, (v) 65 encircled molecules. b. Corresponding $\boldsymbol{<}{\hat{\boldsymbol{L}}}_{\boldsymbol{100}}\boldsymbol{>}$ values extracted from all molecules over time (depicted blue) and their associated fixed targets (depicted red). c. Convergence time and d. Error as a function of the number of molecules for each *L_100_* target (depicted in blue are the values obtained for each target).

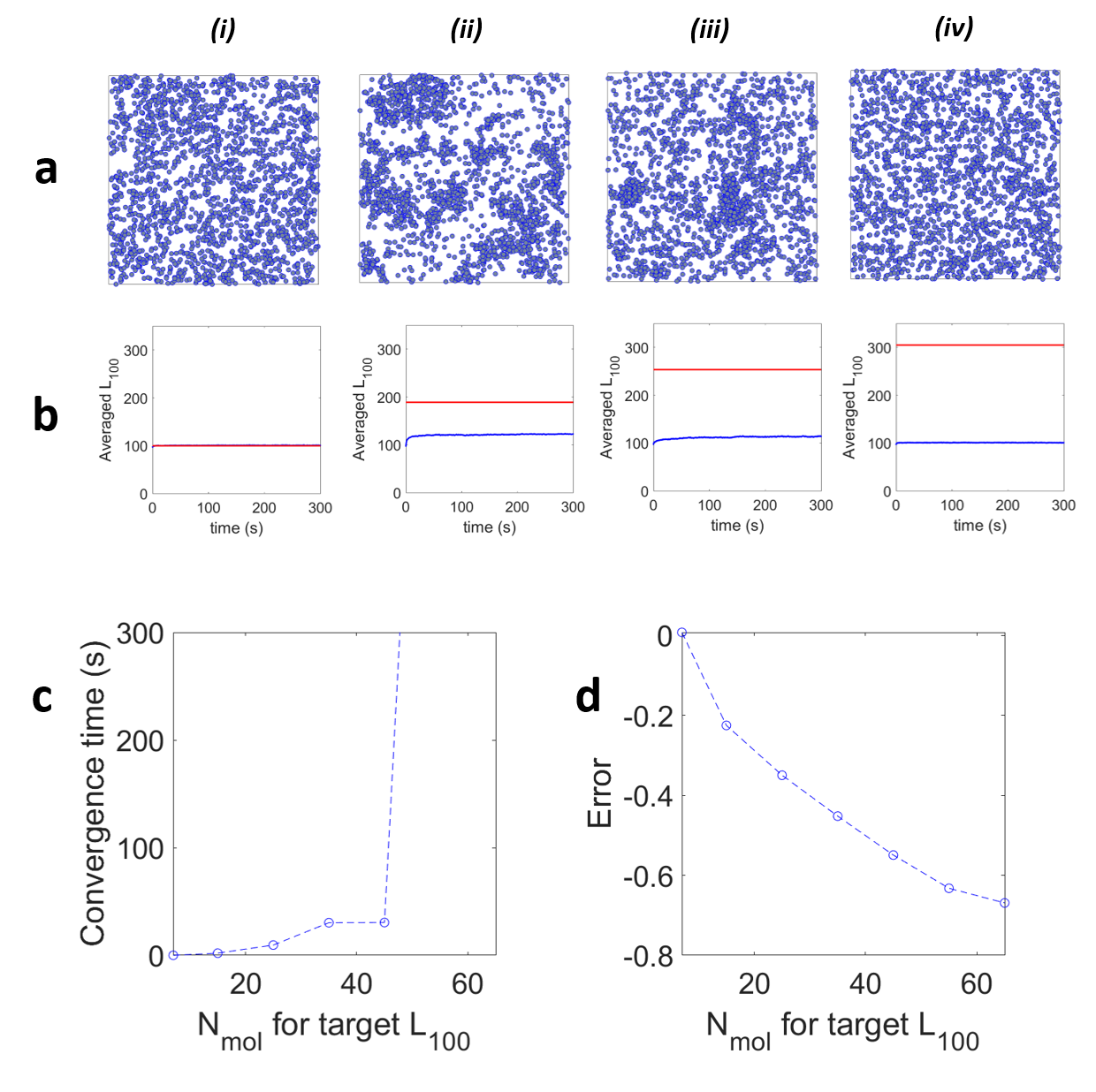

Figure Supplementary 8: Converging scenario with Dmin ~ 0.93 nm.ms^-1,^ (n=10 simulations per condition). a. Examples of the molecular distributions obtained after 5 min simulations with the target values set at (i) 7, (ii) 25, (iii) 45, (iv) 55, (v) 65 encircled molecules. b. Corresponding $\boldsymbol{<}{\hat{\boldsymbol{L}}}_{\boldsymbol{100}}\boldsymbol{>}$ values extracted from all molecules over time (depicted blue) and their associated fixed targets (depicted red). c. Convergence time and d. Error as a function of the number of molecules for each *L_100_* target (depicted in blue are the values obtained for each target).

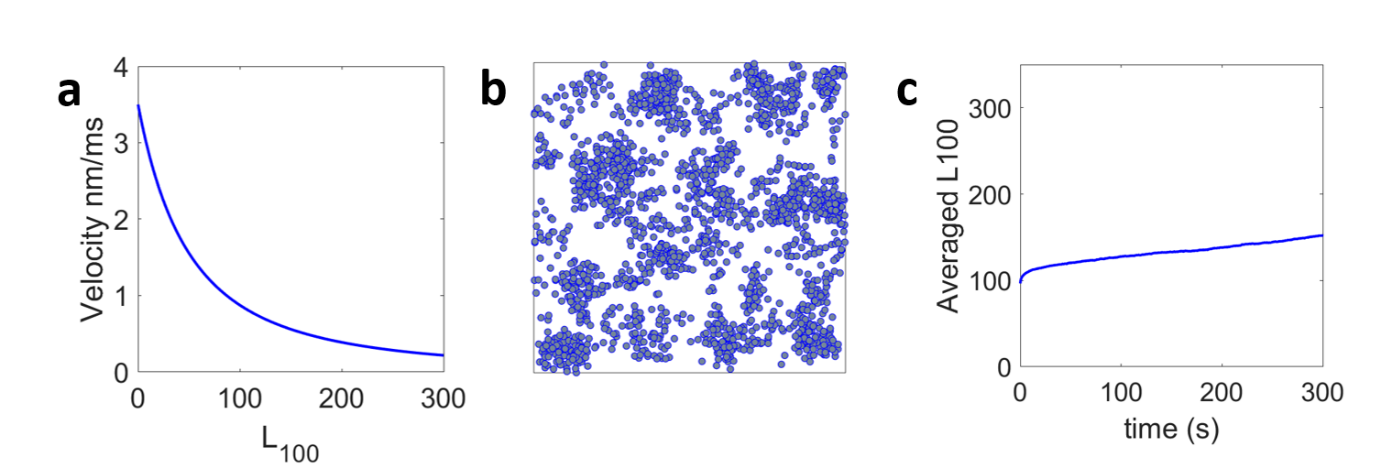

Figure Supplementary 9: Quadratic decay, (n=30 simulations per condition). a. decay used for the displacement calculation of each molecule at each frame. b. Example of the molecular distributions obtained after 5 min. c. Corresponding $\boldsymbol{<}{\hat{\boldsymbol{L}}}_{\boldsymbol{100}}\boldsymbol{>}$ values extracted from all molecules over time (depicted blue).

### Supplementary movies

Supplementary Movie 1: Diffusing molecules over the 5 minutes simulation with 1 frame per sec, sped 10 times, in the Standard Condition. Target value: 41 encircled molecules.

Supplementary Movie 2: Diffusing molecules over the 5 minutes simulation with 1 frame per sec, sped 10 times, in the case of slowly diffusing molecules. Target value: 41 encircled molecules.

Supplementary Movie 3: Diffusing molecules over the 5 minutes simulation with 1 frame per sec, sped 10 times, in the case of fast diffusing molecules. Target value: 41 encircled molecules.

Supplementary Movie 4: Diffusing molecules over the 5 minutes simulation with 1 frame per sec, sped 10 times, with a non-converging displacement profile applied to agent-molecules. Target value: 41 encircled molecules.

Supplementary Movie 5: Diffusing molecules over the 5 minutes simulation with 1 frame per sec, sped 10 times, with a converging displacement profile for which D_min_ > 0 applied to agent-molecules. Target value: 35 encircled molecules.
